# Generation of Gausemycin A-resistant *Staphylococcus aureus*

**DOI:** 10.1101/2022.04.26.489521

**Authors:** Darya V. Poshvina, Diana S. Dilbaryan, Sergey P. Kasyanov, Vera S. Sadykova, Olda A. Lapchinskaya, Eugene A. Rogozhin, Alexey S. Vasilchenko

## Abstract

Gausemycins A and B are the first members of the novel lipoglycopeptides family produced by *Streptomyces roseoflavus* INA-Ac-5812, which showed the ability to fight clinically important Gram-positive bacteria, including methicillin-resistant *Staphylococcus aureus*. However, new antibiotics need to be studied in depth to determine their full potential. In this study, we concentrated our efforts to investigate resistance emerging within *S. aureus* upon gausemycin A application.

Using serial passaging of *S. aureus* FDA209P in increasing concentrations of gausemycin A, we obtained the resistant variant *S. aureus* 5812R which are 80-times more resistant comparing to the origin strain. Moreover, obtained resistance is stable, since 15 passages in a drug-free medium did not restore bacterial susceptibility to gausemycin A.

Elucidating of the differences between resistant and parent strains was concerned antibiotic cross-resistance, structure of bacterial membrane, and response at genetic level.

Susceptibility testing of *S. aureus* 5812R revealed the acquisition of cross-resistance to daptomycin, cefazolin, and tetracycline, while resistance to vancomycin, nisin and ramoplanin absence. The composition of fatty acids constituting the cytoplasmic membrane of *S. aureus* 5812R, was represented by increased content of *anteiso*- branched chain fatty acids, while *iso*-branched chain fatty acids was decreased comparing the origin *S. aureus* FDA209P strain. The relative expression of the *cls* gene catalyzing the synthesis of cardiolipin in the resistant cells was higher compared to the *S. aureus* FDA209P.

## 1. Introduction

Antimicrobial resistance (AMR) possesses a serious global threat of growing concern to human, animal, and environment health. The emergence and rapid spread of pathogenic microorganisms with multidrug resistance (MDR) to antibiotics, observed in recent decades, is especially dangerous [1],[2],[3]. The list of the main Gram-positive pathogens includes methicillin-resistant *S. aureus* (MRSA) and vancomycin-resistant *S. aureus* (VRSA), since they play a leading role in the etiology of a wide range of community-acquired and hospital-acquired human infections [4],[5],[6]. It is important to note that most of the new antibiotics discovered over the past 30 years show activity mainly against Gram-positive bacteria [7], in this regard, antibiotic resistance among Gram-positive bacteria is an alarming phenomenon.

The list of novel antibiotics which are proposed as alternative to conventional ones, contains various antimicrobial peptides (AMPs), including bacteriocins and lipopeptides with a broad spectrum of antimicrobial activity, [8],[9],[10],[11],[12],[13]. Non-ribosomal peptide antibiotics (NRPs) are of considerable interest from the standpoint of therapeutic potential [14]. Glycopeptide vancomycin is a conventional antibiotic with a long clinical history, while lipopeptide daptomycin was approved by the US Food and Drug Administration (FDA) in 2003 for the treatment of skin and soft tissue infections caused by *S. aureus* [15]. Although daptomycin has become available in medical practice relatively recently, in various geographical regions the resistant isolates have been already appeared [16].

Recently the new lipoglycopeptides were revealed among metabolites of actinomycete *Streptomyces roseoflavus* INA-Ac-5812. These lipoglycopeptides were first described in 2016 [17] and then were named as gausemycins A and B [18].

Gausemycin A is cyclic lipoglycopeptide, resembling anionic lipopeptides, but with a completely distinct peptide sequence. The closest structurally related compounds are the depsipeptides (daptomycin, taromycins, and cadazides) and the cyclopeptides (amphomycin, rumycins, and malacidins) [18].

Gausemycin A demonstrated mode of action similar with daptomycin, but had some otherness. Gausemycin A has a pronounced activity against Gram-positive bacteria, including methicillin-resistant *Staphylococcus aureus* (MRSA). In addition, gausemycin A has an inhibitory effect not only on planktonic cells, but also on biofilm-embedded cells [19].

Thus, the above properties of gausemycin A make it as an alternative to existing antibiotics for the treatment of infectious diseases caused by *S. aureus.* However, before the new substance will moving to full preclinical investigations of it is necessary to evaluate its ability to generate antibiotic-resistant forms. These inevitable cycles of antibiotic discovery and resistance not only pose the risk of over-reliance on peptide antibiotics, but also highlight the importance of elucidating emerging resistance mechanisms [20].

In this regard, the aim of this work was to evaluate the ability of ubiquitously used *S. aureus* FDA209P (ATCC 6538P) strain to develop resistance to new lipoglycopeptide gausemycin A, and recognize the phenotypic changes occurring in the *S. aureus* population under the treatment.

## 2. Results

### 2.1 The dynamics of the resistance formation

The first goal of this investigation was to evaluate the dynamics of the resistance development of *S. aureus* FDA209P to gausemycin A in comparison with ramoplanin and daptomycin, which have similar structure and mechanism of action, respectively.

The resistant population of *S. aureus* FDA209P was obtained by multiple serial passages with sub-inhibitory concentrations of gausemycin A. The formation of resistant variant of *S. aureus* FDA209P to gausemycin A occurred within twenty passages (Table 1).

**Table 1.**
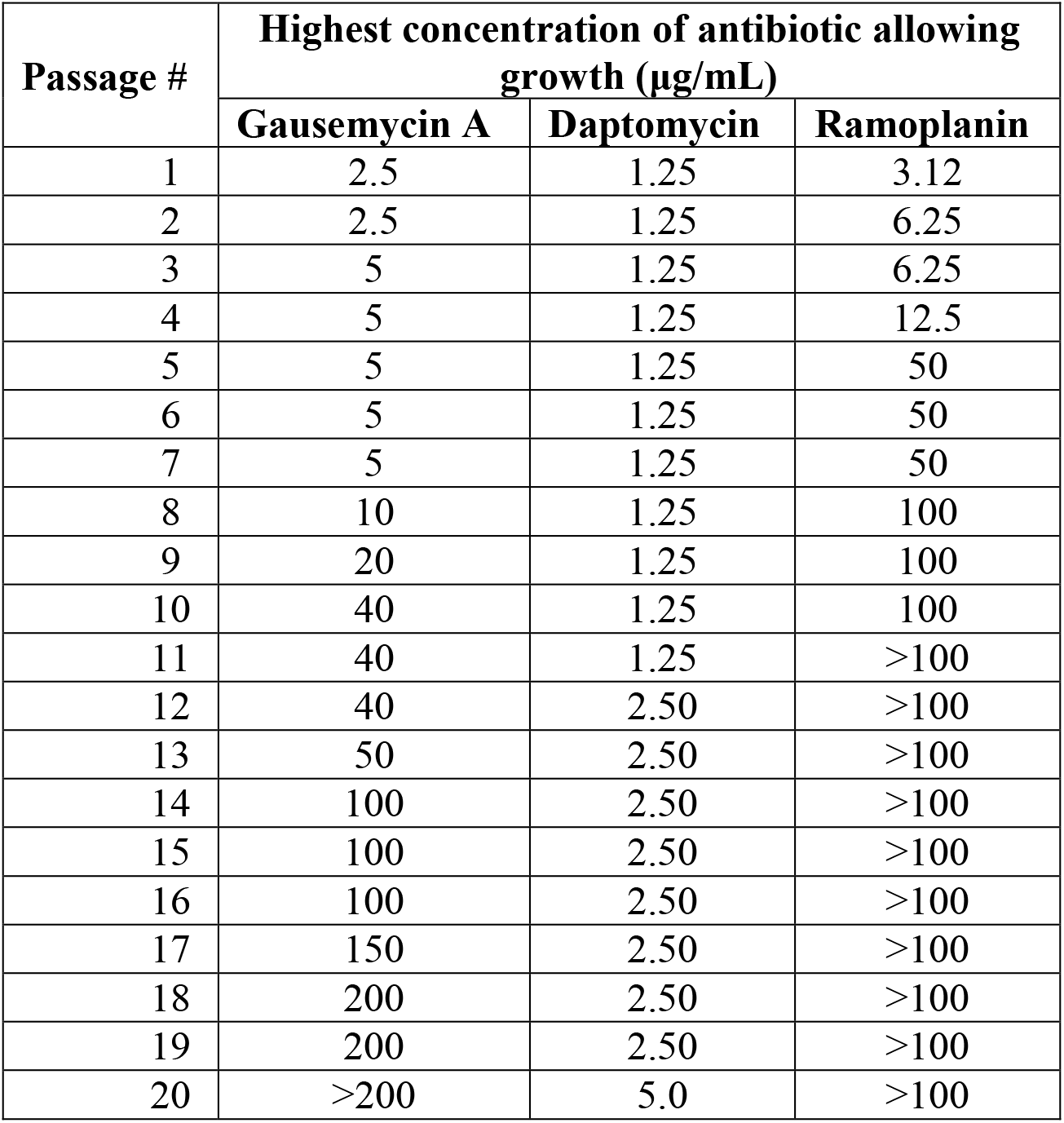
Serial passage of *S. aureus* FDA209P in increasing concentrations of antimicrobial peptides

From the 12th passage with gausemycin A, the bacteria became significantly less sensitive. The obtained clone of *S. aureus* FDA209P at the 13th passage was resistant to the antibiotic concentration in the medium equal to 50 μg/mL.

Adapted population was found to display up to 80-fold increment in their minimum inhibitory concentration (MIC) relative to the wild-type strain. Resistance formation of *S. aureus* FDA209P to ramoplanin occurred within ten passages. Already from the 5th passage of cultivation with ramoplanin, the bacteria became significantly less sensitive to this antibiotic. Eight passages were enough to reach MIC-value for 100 μg/ml. At the same time, the increase in resistance of *S. aureus* FDA209P to daptomycin during a series of passages was less pronounced. Daptomycin’s minimum inhibitory concentration (MIC) was increased from 1.25 μg/mL to 5.0 μg/mL at 20 passage (Table 1).

### 2.2 *Cross-resistant of S. aureus* 5812R

The cross-resistance of obtained *S. aureus* 5812R strains was evaluated against the main class of antibiotics, which target is a Gram-positive bacterial’ barrier structures (cell wall and/or cytoplasmic membrane), with the exception of tetracycline, the target of which is 50S ribosome subunit. It was found, that the most significant difference in susceptibility was acquired to daptomycin, which was 4-fold more potent against *S. aureus* FDA209P comparing to *S. aureus* 5812R. The MIC values for cefazolin and tetracycline were reduced by 2 times for *S. aureus* FDA209P comparing to *S. aureus* 5812R. Surprisingly, but two comparing strains did not differ in their susceptibility to vancomycin, ramoplanin and nisin (Table 2).

**Table 2.**
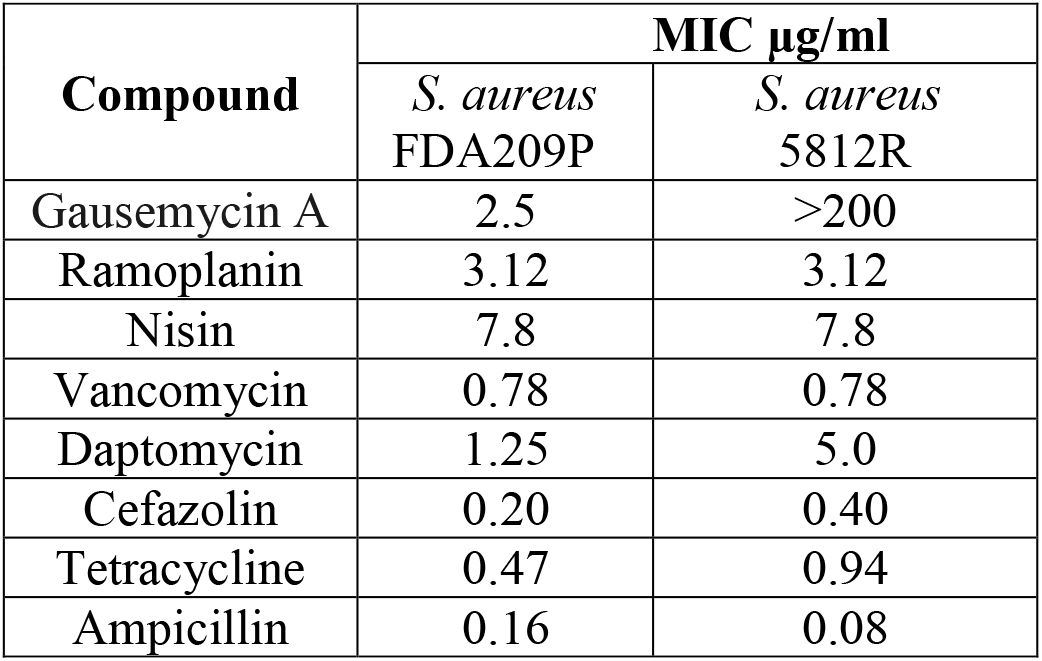
Susceptibility of *S. aureus* FDA209P and *S. aureus* 5812R to various antibiotics

### 2.3 The obtained resistance is not reversible

The obtained gausemycin A resistant clone retained signs of stability after multiple (17 times) passages on the gausemycin A-free media. The effect of gausemycin A on the growth dynamics of the *S.aureus* FDA209P strain was monitored spectrophotometrically in 96-well microtiter plate, which allowed simultaneous cultivation and analysis of bacterial growth in real time. The resistant strain demonstrated comparable with origin strain growth rate, but decreased population density at the end-point of measurement (Figure 1).

**Figure 1.**
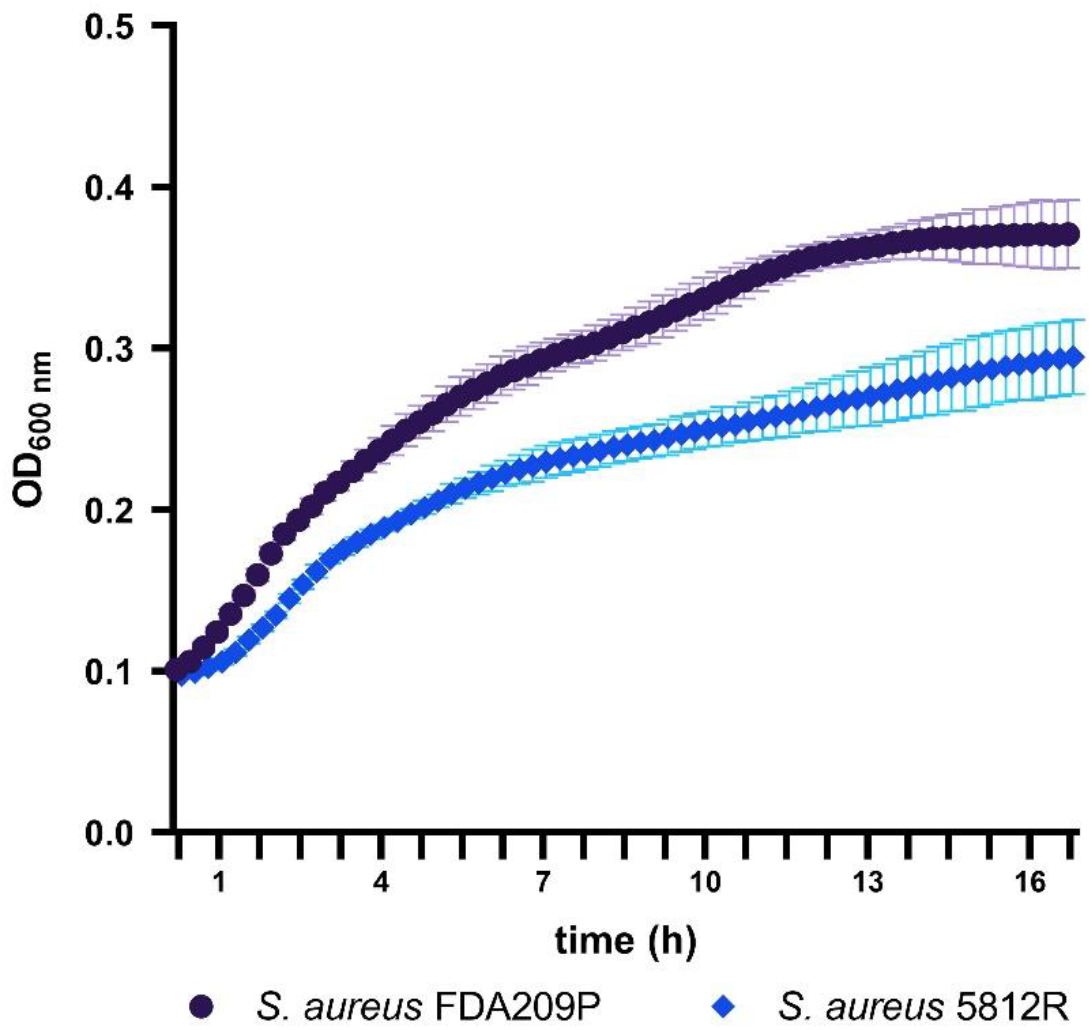
Growth kinetics of the *S. aureus* FDA209P and *S. aureus* 5812R on Muller-Hinton medium without gausemycin A.

### 2.4 *Fatty acid profiles of S. aureus* FDA209P *and S. aureus* 5812R

*S. aureus* demonstrates considerable membrane lipid plasticity in response to different growth environments, which is of potential relevance to response and resistance to various antimicrobial agents [25]. As shown in Table 3, a total of 26 fatty acids were identified in the membrane of *S. aureus.* The straight chain fatty acids (SCFAs) and the branched chain fatty acids (BCFAs), including *iso*- and *anteiso*-BCFAs for *S. aureus* FDA209P constituted 19.83%, 18.06% and 57.76% of all membrane fatty acids, respectively (Figure 2). For *S. aureus* 5812R the relative proportions of *anteiso*-BCFAs increased to 64.4%, whereas iso-BCFAs decreased to 14.64%. The relative proportions of saturated fatty acids changed slightly from 19.83% for the *S. aureus* FDA209P to 19.26% for the *S. aureus* 5812R. The increase of *anteiso*-BCFAs was mostly attributed to the increased level of *anteiso*-C15:0 and *anteiso*-C17:0, which proportions increased from 33.41% and 16.84% to 36.75% and 19.99%, respectively, in clones of *S. aureus* FDA209P resistant to gausemycin A. Comparatively, the decrease of *iso*-BCFAs was mostly attributed to *iso*-C15:0, *iso*-C17:0 and *iso*-C19:0. Our finding suggests that *S. aureus* 5812R responded effectively to the toxic stress of gausemycin A through changing the proportions of BCFAs.

**Figure 2.**
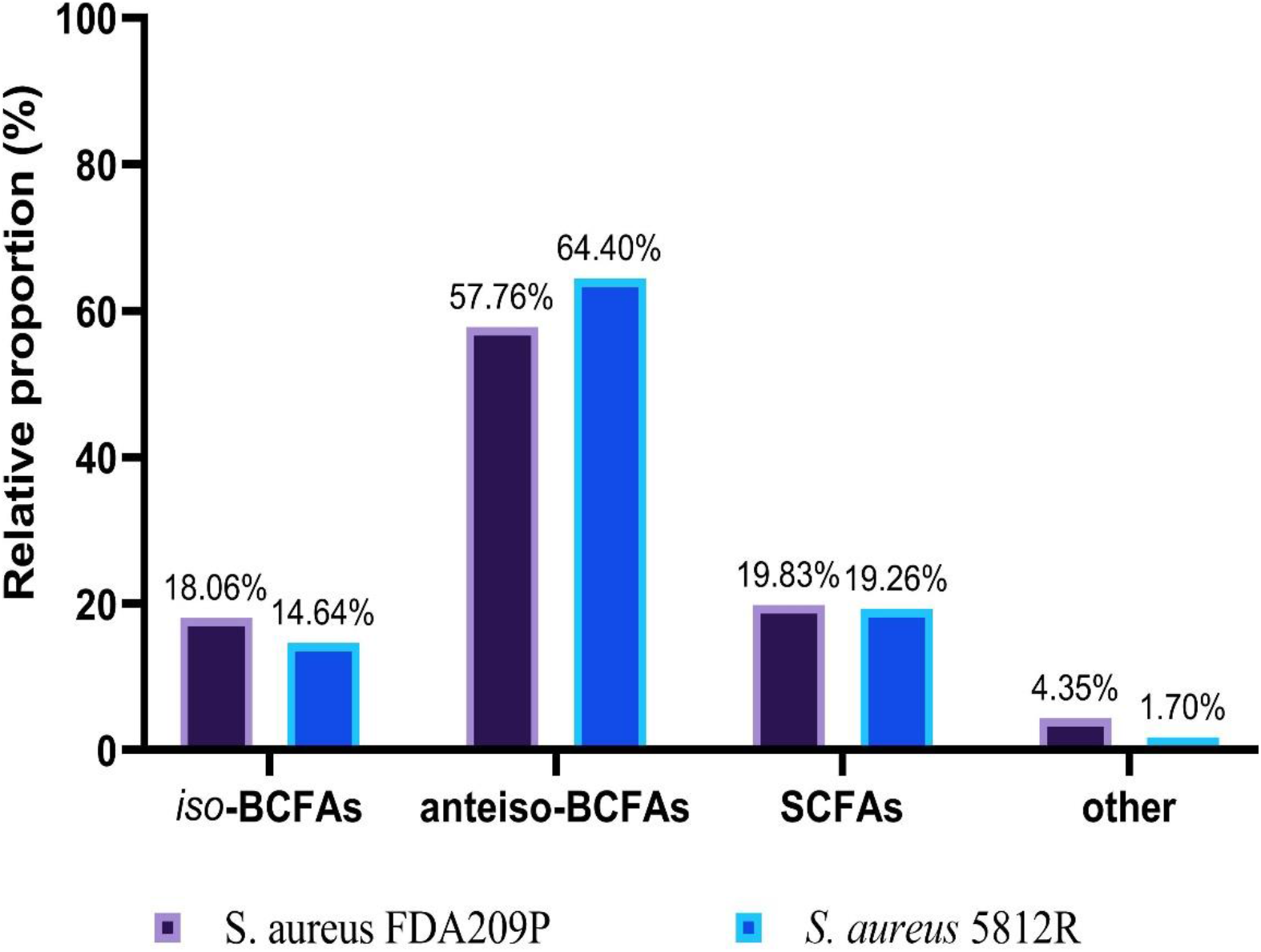
Fatty acid composition of cytoplasmic membrane *S. aureus* FDA209P and *S. aureus* 5812R

**Table 3.**
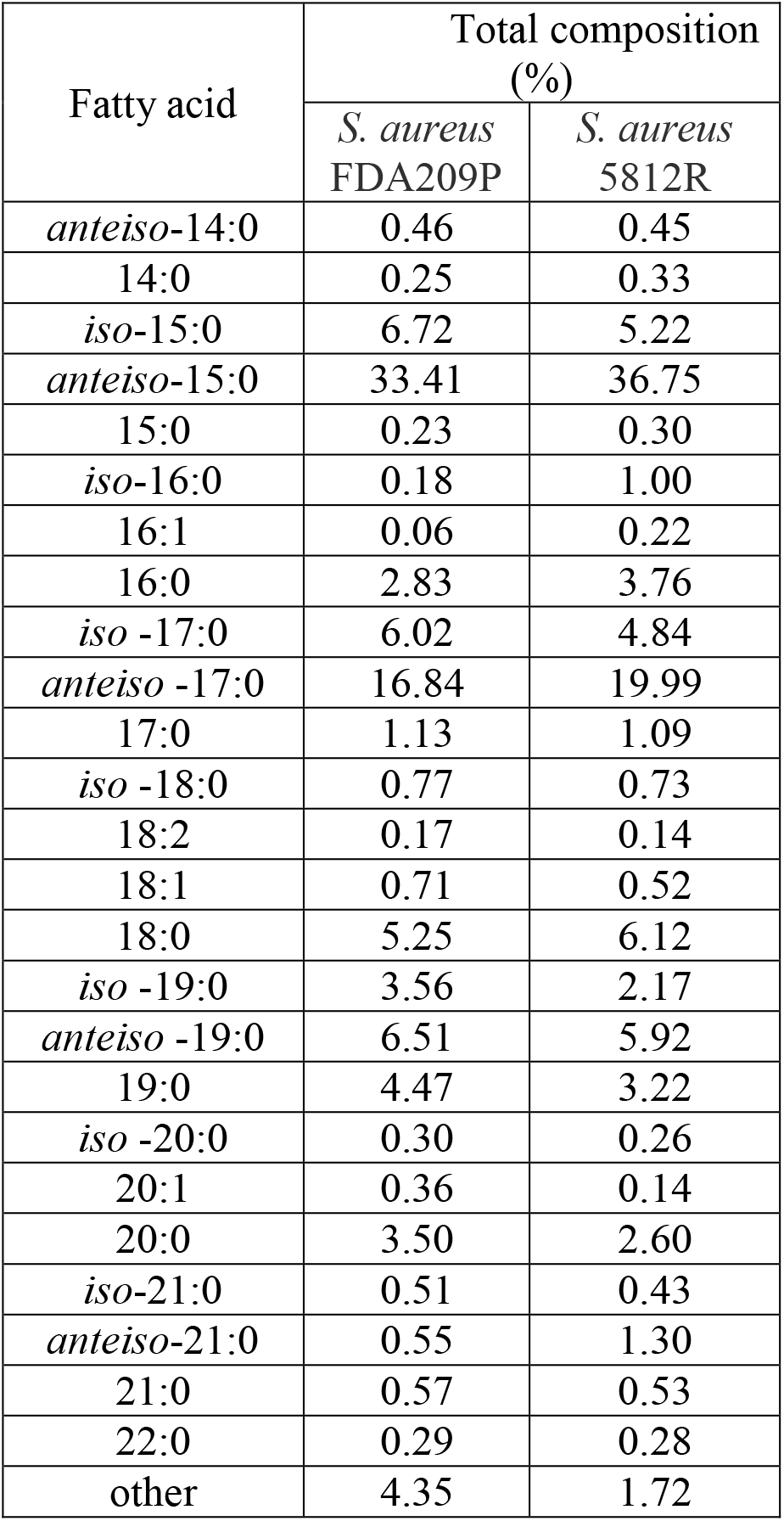
Membrane fatty acid composition of stationary stage cells of *S. aureus*

| Fatty acid | Total composition (%) |  |
| --- | --- | --- |
|  | *S. aureus* FDA209P | *S. aureus* 5812R |
| *anteiso*-14:0 | 0.46 | 0.45 |
| 14:0 | 0.25 | 0.33 |
| *iso*-15:0 | 6.72 | 5.22 |
| *anteiso*-15:0 | 33.41 | 36.75 |
| 15:0 | 0.23 | 0.30 |
| *iso*-16:0 | 0.18 | 1.00 |
| 16:1 | 0.06 | 0.22 |
| 16:0 | 2.83 | 3.76 |
| *iso* -17:0 | 6.02 | 4.84 |
| *anteiso* -17:0 | 16.84 | 19.99 |
| 17:0 | 1.13 | 1.09 |
| *iso* -18:0 | 0.77 | 0.73 |
| 18:2 | 0.17 | 0.14 |
| 18:1 | 0.71 | 0.52 |
| 18:0 | 5.25 | 6.12 |
| *iso* -19:0 | 3.56 | 2.17 |
| *anteiso* -19:0 | 6.51 | 5.92 |
| 19:0 | 4.47 | 3.22 |
| *iso* -20:0 | 0.30 | 0.26 |
| 20:1 | 0.36 | 0.14 |
| 20:0 | 3.50 | 2.60 |
| *iso*-21:0 | 0.51 | 0.43 |
| *anteiso*-21:0 | 0.55 | 1.30 |
| 21:0 | 0.57 | 0.53 |
| 22:0 | 0.29 | 0.28 |
| other | 4.35 | 1.72 |

### 3.5 *Comparison of cardiolipin synthase (cls) gene expression between S. aureus* FDA209P *and S. aureus* 5812R

Cardiolipin plays a critical role in bacterial physiology, especially for cytokinesis and antibiotic resistance [26]. The synthesis of cardiolipin in bacteria is catalyzed by cardiolipin synthase (Cls), which provides condensation of two phosphatidylglycerol molecules to yield cardiolipin and glycerol.

We analyzed the expression of the *cls* (cardiolipin synthase) gene in both strains in comparing manner.

It was found significantly higher transcriptional level of *cls* gene in the cells of resistant *S. aureus* 5812R (Figure 3). However, *cls* transcription have been decreased with time in both stains in the same manner.

**Figure 3.**
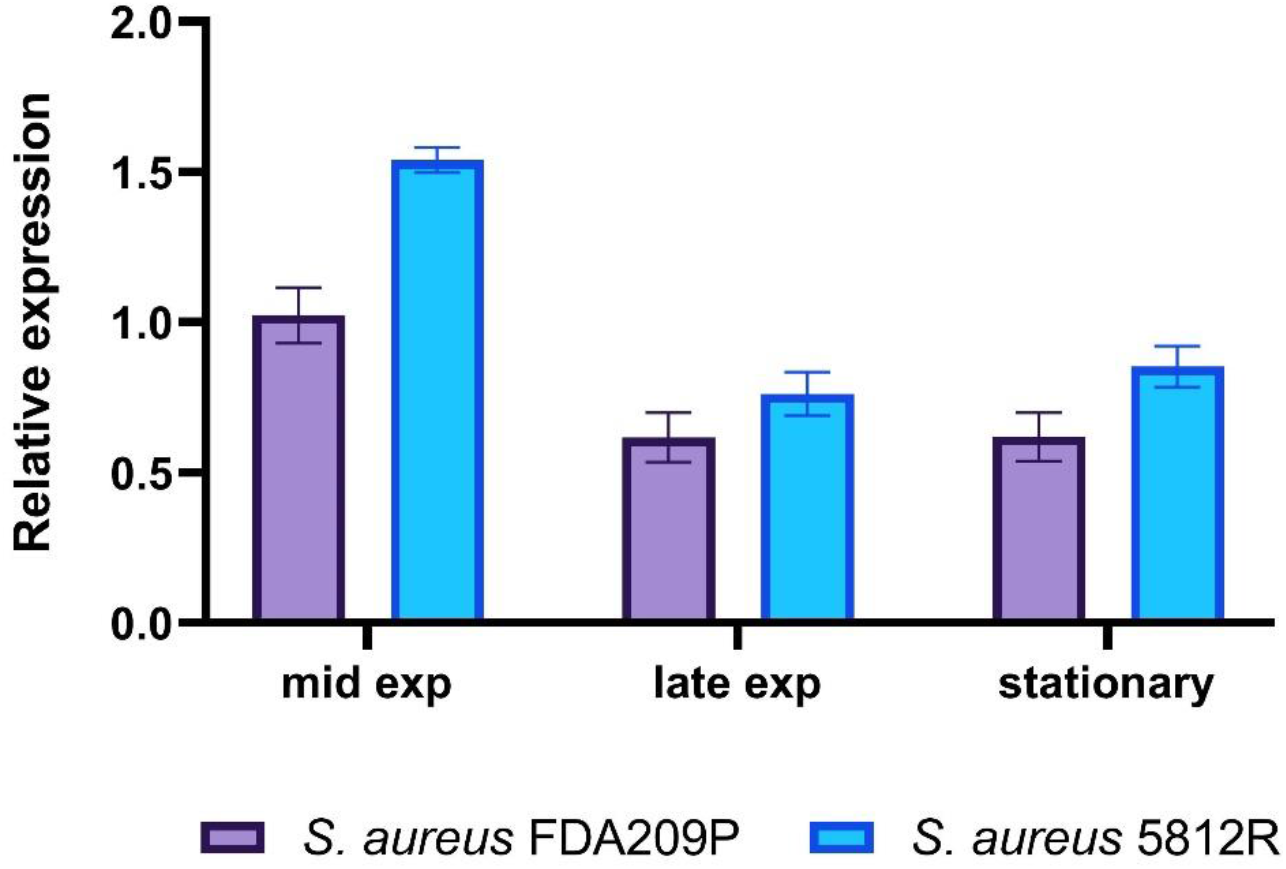
Expression analysis of gene responsible for the synthesis of cardiolipin (*cls*). Transcript levels of the analyzed genes were determined by RT-qPCR in relation to *gyrB* expression. Mid Exp, mid-exponential growth phase; Late Exp, late exponential growth phase; Stationary, stationary growth phase.

### 2.6 Membrane integrity assay

Our previous study showed that the mechanism of action of gausemycin A (5812-A/C) is aimed at disrupting the structural integrity of cell membranes [19]. Method for detecting membrane-disturbing action is simple and based on the detection of quenching of the SYTO9 green fluorescence, when red-fluorescent dye propidium iodide influx through disordered bacterial membrane. If the resistance of staphylococci is associated with a change in the membrane structures of the cells, then there will be no quenching of SYTO9 fluorescence. It was shown that the addition of gausemycin A into the reaction medium quenched SYTO9 fluorescence in dose-dependent manner (Figure 4), and complete quenching was achieved at a 50μg/ml. In contrary, it was unable to achieve reducing in SYTO9 fluorescence with *S. aureus* 5812R which is resistant to gausemycin A. Collectively, obtained data is evidence what resistance to gausemycin A is related to membrane modifications, and contrary, this fact is an additional argument in favor of membrane permeabilizing mechanism of gausemycin A’s action.

**Figure 4.**
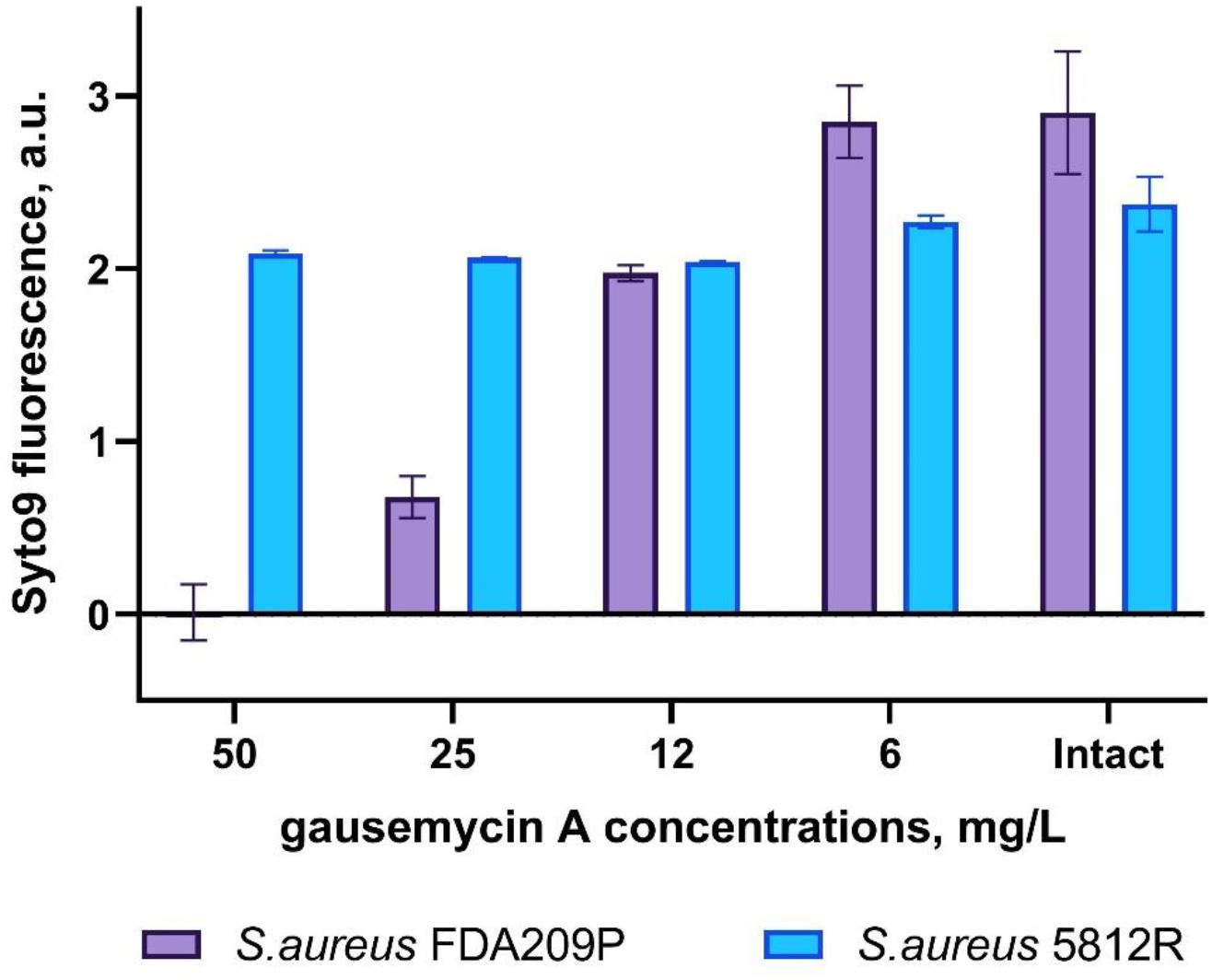
Evaluation of membrane disturbance of *S. aureus* FDA209P and *S. aureus* 5812R using Live/Dead fluorescent dyes. If bacterial membranes are permeabilized, propidium iodide penetrates into the cell. What follows is SYTO 9 getting displaced from nucleic acids, which leads to a decrease in fluorescence intensity in the green region (505–540 nm) of the spectrum. The data presented as intensity of Syto9 fluorescence at the end-point of measurement (10 min).

## 3. Discussion

Given the continued high interest in antimicrobial peptides as potential therapeutics for the treatment of bacterial infections, it is especially important to assess the risk of resistance development and study its mechanisms. Generation of a *S. aureus* FDA209P strain with reduced susceptibility to gausemycin A provides a model system to achieve greater insight into the evolution of resistance mechanisms in Gram-positive bacteria against peptide antibiotics. The present work is the only second evidence of bacterial resistance against lipoglycopeptides after the Schmidt’s study of ramoplanin resistant *S.aureus* [27].

Gausemycins are the cyclic lipoglycopeptides, resembling anionic charge but with a completely distinct peptide sequence. The barrier structures of Gram-positive bacteria represented by peptidoglycan shell and cytoplasmic membrane, which are play a fundamental role in the functioning and survival of bacterial cells. We have taken daptomycin and ramoplanin to compare the features of the gausemycin A resistance formation against two antibiotics which having certain similarities with gausemycin A. Earlier, we showed that gausemycin A has a mode of action similar to that of membrane-active lipopeptide daptomycin [19], while structurally more resemble lipoglycopeptide ramoplanin [18].

Obtained results indicate that gausemycin A differs from these compared antibiotics. *S. aureus* FDA209P acquired significant resistance (16-fold) to ramoplanin at fifth passage, while resistance to daptomycin did not exceed initial level more than 4-fold by the 20th passage. According to the obtained results, gausemycin A has an intermediate position, since a 16 times more resistant clone of *S. aureus* was formed by the 10th passage.

Thus, development of resistance to gausemycin A in comparison with antibiotics which are similar in structure and/or mechanism of action was much faster than to daptomycin, but much slower than to ramoplanin.

In favor of the fact that the mechanism of the antimicrobial action of gausemycin A is unique, the results of the detection of cross-resistance of the *S. aureus* 5812R strain testify. So, resistance to ramoplanin, nisin and vancomycin has not been identified. These peptide antibiotics unites together by the same target within cells. For example, vancomycin is a cationic glycopeptide antibiotic that kills bacteria by binding to the C-teminal D-Ala–D-Ala residues of the peptidoglycan precursor lipid II, that prevent the use of the precursor for cell wall synthesis [28]. Nisin has a membrane-destroying effect upon binding to intracellular receptor, which is lipid II [29]. The pyrophosphate pocket of lipid II is a target for ramoplanin too [30].

Thus, decreasing the amount of lipid II would be a one of the existing mechanisms for generation of bacterial resistance. Giving the fact that nisin, ramoplanin and vancomycin are cationic molecules, while gausemycin A has anionic in nature, so it is obvious that resistance to gausemycin A is not formed due to peptidoglycan modification.

The next task was to find out what is the cause of resistance? First of all, it was necessary to evaluate the structural changes that occurred in the cytoplasmic membrane, since it is the target of gausemycin A.

Among the adaptive changes in the chemical composition of the cell envelope that have been well documented is a change in cytoplasmic membrane phospholipid composition. Importantly, the majority of *S. aureus* membrane lipids are comprised of negatively charged phospholipids: phosphatidylglycerol (PG) and cardiolipin (CL) [31]. The known position of cardiolipin is due to the presence of a molecular structure. Having 4 fatty acid residues and 2 phosphoric acid residues, cardiolipin has the function of a proton and electron donor in the membrane, regulates membrane permeability, and performs a stabilizing function [32]. Under conditions of stress, such as unfavorable growth conditions or cell-wall acting antibiotics, cardiolipin can accumulate up to ~25-30 % of membrane phospholipid [33].

In bacteria, cardiolipin synthases (Cls) are the critical enzymes for the synthesis of CL, often using two molecules of PG as substrate. Analysis of the *cls* gene expression of *S. aureus* FDA209P and *S. aureus* 5812R showed that the development of resistance to gausemycin A is accompanied by an increase in the relative expression of the *cls* gene for *S. aureus* 5812R. It may be one of the factors contributing to the increase in the resistance of the *S. aureus* 5812R to gausemycin A. As for cardiolipin, gausemycin A is not the only antibiotic whose activity is affected by the content of this phospholipid. Enhanced levels of cardiolipin are associated with daptomycin resistance in *Enterococcus faecalis* and *Enterococcus faecium* or *Staphylococcus aureus* [34].

The bacterial membrane equilibrium between rigidity and fluidity is driven by multiple factors that impact membrane order (water, metal ions, proteins, carotenoids, staphyloxanthin, etc.), but it is primarily related to lipid packing within the membrane, which is determined by the structure and composition of phospholipid fatty acids [35]. In the case of *S. aureus*, bacterial membranes mainly consist of straight-chain and branched-chain saturated fatty acids. Straight-chain fatty acids pack together to produce a bilayer with low permeability properties, while branched-chain *iso* or *anteiso* methyl species promote a more fluid membrane structure [36].

Changes in the ratio of BCFAs to SCFAs to regulate the cytoplasmic membranes fluidity of Gram-positive bacteria is one of the most common adaptation mechanisms [37]. The development of resistance of *S. aureus* 5812R to the studied lipoglycopeptide was accompanied by an increase in the relative proportions of *anteiso*-BCFAs in cell membranes at the same time by a decrease in the levels of *iso*-BCFAs. The percentage of saturated fatty acids in the cell membrane of the resistant strain decreased slightly compared to the wild type. This result was inconsistent with previous studies showing that *S. aureus*, reduced the fluidity as a result of an increased level of SCFAs in the unfavorable conditions. For example, *S. aureus* responded to higher concentrations of daptomycin and vancomycin by lowering the biosynthesis of *iso* and *anteiso* branched-chain fatty acids while increasing the levels of saturated fatty acids [35], [38].

However, such alterations in membrane fatty acid composition were consistent with some previous works showing the proportion of BCFAs would increase for some Gram-positive bacteria in undesired environmental conditions. Sun et al. [39] reported that fatty acids with *anteiso* branching are more effective at fluidizing the membrane than fatty acid with *iso* branching and there is a correlation between an increase in the proportion of *anteiso* fatty acids as an adaptive response to reduced temperatures that is especially important for the survival of cold tolerant pathogens like *Listeria monocytogenes*. Changes in the ratio of BCFAs to SCFAs to regulate the cytoplasmic membranes fluidity of Gram-positive bacteria is one of the most common adaptation mechanisms

The findings suggest that bacteria can potentially develop possible adaptive mechanisms to create tolerance for antibacterial activity of gausemycin A at a low level. Further study of gausemycin A-resistant *S. aureus* 209P will provide insight into the molecular mechanisms of gausemycin A resistance in *S. aureus* and may lead to strategies to prevent antimicrobial resistance during clinical use.

## 4. Conclusion

In this study we showed the ability of *S.aureus* 209P to form gausemycin A resistant phenotype very quickly. However, revealed phenomenon does not mean that this peptide antibiotic has no prospects as a therapeutic agent. Ramoplanin showed an even more pronounced ability to form a resistant *S. aureus* phenotype in this comparative experiment. In addition, *S. aureus* 5812R did not form cross-resistance to antibiotics that have other cellular targets, which allows considering gauzemycin A in combination therapy.

Another kind of obtained result, concerns the mechanism of gausemycin A action. We demonstrated that cross-resistance of *S. aureus* 5812R to daptomycin, structural changes in bacterial cytoplasmic membrane and increased expression of the *cls* gene, collectively once again demonstrates the similarity between gausemycin A and daptomycin.

## 5. Materials and methods

### 5.1 Bacterial culture

*Staphylococcus aureus* FDA209P (MSSA, methicillin-susceptible) sensitive to gausemycin A and its resistant variant (*S. aureus* 5812R) resulting from a series of passages used in this study.

### 5.2 Serial passage experiments

To generate gausemycin A resistant variant, a step pressure method was employed. Isolated colonies of *S. aureus* FDA209P were inoculated into Mueller-Hinton II broth (CaCl_2_ 50mg/L) containing gausemycin A. The cultures were incubated at 37 °C with aeration for 24 h. After incubation, bacterial cells growing at the highest concentration of gauzemycin A (below the MIC) were harvested and used for next passage. The process was repeated for 20 passages. The dynamic of the formation of resistance to gausemycin A was evaluated in comparison with daptomycin and ramoplanin.

### 5.3 Determination of minimum inhibitory concentrations (MIC)

Determination of minimal inhibitory concentration (MIC) was performed as described in method [21], following the Clinical and Laboratory Standards Institute (CLSI) guidelines for broth microdilution MIC assays. Bacteria were incubated in a 96-well microtiter plate (Eppendorf, Germany) containing 90 μl of inoculum prepared in growth media at 10^6^ CFU/ml with 10 μl of 2-fold dilutions of the antibiotics. The results were evaluated after 24 h of cultivation at 37 °C. The dynamics of bacterial growth were assessed by scanning and plotting the absorbance data at 600 nm obtained by spectrophotometer (Multiscan GO, Thermo Fisher Scientific, United States). Antimicrobial susceptibility to daptomycin, vancomycin, ramoplanin, nisin, cefazolin, ampicillin and tetracycline was tested for gausemycin A-susceptible and resistant strains. Stock solutions (2 mg/ml) were prepared in ddH_2_O and filter sterilized. Broth microdilution assay was performed in a 96-well plate as described above. All experiments were performed in triplicate.

### 5.4 Resistance stability testing

Overnight culture of *S. aureus* 5812R was sub-cultured in gauzemycin A-free Muller-Hinton broth at a cell density of 10^6^ CFU/mL and incubated overnight at 37°C with shaking. The obtained culture was evaluated on susceptibility to gausemycin A in MIC assay as described previously. Passages without selection continued for 17 times.

### 5.5 Membrane lipid extraction, fractioning and methylation

Fatty acid composition was determined using biomass of *S. aureus* FDA209P and *S. aureus* 5812R. Bacteria grown to stationary phase in Muller-Hinton medium with CaCl_2_ at 37°C with shaking until reaching an OD 620 nm 0.5 (10^9^ CFU/mL). Cells were harvested by centrifugation and washed two times with phosphate-buffered saline and lyophilized.

Lipid extraction was performed according to Folch method [22]. Chloroform:methanol (2:1, v/v) was added to each sample according to the original Folch procedure allowing for a CHCl_3_:MeOH:H_2_O ratio of 8:4:3 (v/v/v). Briefly, ice-cold methanol and chloroform were added to the sample. The suspension was vortexed occasionally to bring about physical mixing and the sample was incubated on ice for 30 min. After the addition of water, which was used to separate the aqueous and organic layers, the suspension was incubated on ice for an additional 10 min. Samples were centrifuged at 2000 rpm for 5 min at 4 °C. The lower phase (organic) layer was transferred to a new tube. The aqueous layer was re-extracted with 1 mL of 2:1, v/v chloroform/methanol. The chloroform layers were combined for analysis. The aqueous layer was centrifuged at 2000 rpm for 5 min at 4 °C and collected for metabolite analysis. Extracts were then dried under nitrogen.

Fatty acid methyl esters (FAME) were obtained by the treatment of the total lipids with 2 % H_2_SO_4_ in MeOH in a screw-capped vial (2h, 80°C) under Argon and purified by thin layer chromatography development in benzene [23]. The GC analysis of FAME was carried out on a Shimadzu GC-2010 chromatograph (Kyoto, Japan) with a flame ionization detector on an Equity-5 (Supelco, Bellefonte, PA, USA) capillary column (30 m/0.25 mm ID, the phase thickness 0.25 lm) at 160 °C with a 2 °C/min ramp to 240 °C that was held for 20 min. Injector and detector temperatures were 250 °C. FAME were identified by GC–MS using a Shimadzu GCMS-QP5050A instrument (Kyoto, Japan) (electron impact at 70 eV) with a SPB-5 (Supelco, Bellefonte, PA, USA) capillary column (30 m /0.25 mm ID). The GC–MS analysis of FAME was performed at 160 °C with a 2 °C/min ramp to 240 °C that was held for 20 min. Injector and detector temperatures were 250 °C. Spectra were compared with the NIST library and FA mass spectra archive (The AOCS Lipid Library 2015).

### 5.6 RNA extraction and real-time quantitative PCR analysis

Each isolate was grown in Mueller Hinton Broth (MHB) in the absence of gausemycin A. The overnight cultures were diluted and as a resulted in starting number of the cells approximately 10^6^ CFU/ml in each culture. Cells were then harvested from the *S. aureus* FDA209P and *S. aureus* 5812R cultures grown to mid exponential (after 4.5 and 6.5 h, respectively), late exponential (after 8 h and 10 h, respectively) and stationary (after 12 h and 14 h, respectively) phase.

Total RNA was isolated using the RNeasy Mini kit (Qiagen, USA) according to the manufacturer’s instructions. Total RNA was further treated with DNase I (New England Biolabs, USA) followed by the RNeasy MinElute Cleanup kit (Qiagen, USA) according to the manufacturer’s instructions. RNA was quantified using Qubit 4.0 (Thermo Fisher Scientific, USA) and the quality of the RNA extracted was assessed by TapeStation 4150 (Agilent Technologies, USA). Samples with preserved 16S and 23S peaks and RIN values >8 were selected for gene expression analyses. Then, using the reversed transcription reaction, complementary DNA was obtained using the iScript reversed transcription supermix for RT-qPCR reagent (Bio-Rad, USA) in accordance with the manufacturer’s protocol. Then quantitative PCR was carried out with the obtained cDNA using SsoAdvanced Universal SYBR Green Supermix reagent (Bio-Rad, USA). Each reaction mix with a volume of 20 μl was prepared with 300 nM each primer (final concentration) and 20 ng of template RNA. A LightCycler96 Real-Time PCR detection system (Roche, Switzerland) was used for the measurements using a protocol with the following thermal cycling conditions: denaturation at 95 °C for 1 min, followed by 40 cycles of denaturation at 95 °C for 10 s and annealing/extension at 60 °C for 15 s. After the last amplification cycle, a melting curve analysis was carried out by heating from 65 to 95 °C in increments of 0.5 °C /s. Negative controls (without template or reverse transcriptase enzyme) were included in each run.

The design of oligonucleotide primer sequences for analysis of the expression of the gene responsible for the synthesis of cardiolipin (*cls*) was carried out using the online service Integrated DNA technologies (https://www.idtdna.com/Primerquest/Home/Index) using sequences of a given gene presented in the GenBank database. Fold changes in the expression levels of the investigated genes were normalized in relation to the levels of *gyr*B mRNA (Table S1).

The relative changes in gene expression were quantified using the Pfaffl method [24]: gene expression ratio = (E target) 1Ct target (control - sample) / (E reference) 1Ct reference (control - sample), where E target is the amplification efficiency of target (gene of interest), E reference is the amplification efficiency of reference (*gyrB*), Ct is the point at which the fluorescence rises above the background fluorescence, 1Ct target is the Ct deviation of the control minus the sample of the target gene transcript, and 1Ct reference is the Ct deviation of the control minus the sample of the reference gene transcript.

## Acknowledgments

This work was supported by the Russian Science Foundation (project number 21-74-00123).

